# Combined Scattering, Interferometric and Fluorescence Oblique Illumination for Live Cell Nanoscale Imaging

**DOI:** 10.1101/2022.07.22.501104

**Authors:** Yujie Zheng, Yean Jin Lim, Hanqi Lin, Tienan Xu, Carmen Longbottom, Viviane Delghingaro-Augusto, Yee Lin Thong, Christopher R. Parish, Elizabeth E. Gardiner, Woei Ming Lee

**Author notes:** co-first authors (Yujie Zheng, Yean Jin Lim). **Address for correspondence** Dr W M Lee, The Australian National University, Canberra ACT 2601, Australia.

## Abstract

To determine the molecular and/or mechanical basis of cell migration using live cell imaging tools, it is necessary to correlate multiple 3D spatiotemporal events simultaneously. Fluorescence nanoscopy and label free nanoscale imaging can complement each other by providing both molecular specificity and structural dynamics of sub-cellular structure. In doing so, a combined imaging system would permit quantitative 3D spatial temporal detail of individual cellular components. In this paper, we empirically determined a series of optimal azimuthal scanning angles and rotating beam to achieve simultaneous and label-free nanoscale and fluorescence imaging. Label-free nanoscale imaging here refers to interferometric, brightfield (BF) and darkfield (DF) rotating coherence scattering (ROCS) microscopy, while fluorescence refers to high inclined Laminated Oblique (HiLO) and total internal reflection fluorescence (TIRF) imaging. The combined capabilities of interferometric, scattering and fluorescence imaging enables (1) the identification of molecular targets (substrate or organelle), (2) quantification of 3D cell morphodynamics, and (3) tracking of intracellular organelles in 3D. This combined imaging tool was then used to characterize migrating platelets and adherent endothelial cells, both critical to the process of infection and wound healing. The combined imaging results of over ∼1000 platelets, suggested that serum albumin (bovine) was necessary for platelets to migrate and scavenge fibrin/fibrinogen. Furthermore, we determine new asynchronous membrane fluctuations between the leading and rear edge of a migrating platelet. We further demonstrated that interferometric imaging permitted the quantification of mitochondria dynamics on lung microvascular cells (HMVEC). Our data suggests that axial displacement of mitochondria is minimized when it is closer to the nucleus or the leading edge of a cell membrane that exhibits retrograde motion. Taken together, this combined imaging platform has proven to quantify multiple spatial temporal events of a migrating cell, that will undoubtedly open ways to new quantitative correlative nanoscale live cell imaging.

## Introduction

Cells direct their movement through the coordination of extra- and intracellular nanoscale movements to fulfil their biological function in immune surveillance ^1^, wound healing ^2, 3^, clot formation ^4^. Live cell imaging is an excellent method to obtain quantitative data molecular and structural events behind cell migration because of high spatial resolution with sub-second temporal resolution ^5, 6^. The core purpose of all cell migration imaging studies is to correlate multiple spatiotemporal cell dynamics at the same time. The spatiotemporal dynamics include (1) structural remodeling of cell cytoskeleton (actin, tubulin), (2) movement of intracellular components (nucleus, mitochondria, endoplasmic reticulum and Golgi), (3) activating transmembrane proteins (4) surrounding mechanical and molecular cues (adhesive molecules ^3, 7^, external fluid stimulus ^8, 9^ and the extracellular matrix - ECM ^10^. Cells must address these four factors that influence their migratory behavior to initiate the formation of membrane protrusions ^8, 11^. The use of multiplex fluorescent labeling strategies ^12^ are constantly being optimized to visualize molecular and structural components of a migrating cell to identify specific molecular engines (Rho GTPases, Wasp/Scar/Wave families and ARP2/3 complex) that drives structural changes in adherent cell before it migrates. But to achieve 3D quantitative imaging with just fluorescence labeling is not trivial because of the inherent risks of photobleaching and phototoxicity ^13^.

Nanoscale label free imaging such as rotating optical coherent scattering (ROCS) has emerged as an important tool to avoid photobleaching and achieve 3D quantitative imaging^14^. The current hallmark of ROCS lies in the ability to extract nanoscale scattering that results in either inverted intensity (brightfield-BF) or bright intensity (darkfield-DF), all of which occurs at > 50° oblique angles^15-17^. While BF and DF ROCS obtains high contrast images based on scattered light from the sample, ROCS does not contain axial positional information of the sample. That said, elastic scattered light from nanoparticles and biological molecules ^18^ do not degrade or saturate over time ^18-20^. However, label free imaging does not provide the molecular identity of molecules and cellular structures. Lately, interferometric signals has been instrumental at tracking axial positions of purified proteins ^19-21^ and membrane protrusions ^22^. Lately, the combinations of ROCS with total internal reflection fluorescence microscopy-TIRF ^17^ as well as interference scattering (iSCAT) with fluorescence confocal ^15^ have begun to play an important role for nanoscopic correlative imaging. The complementary elements of fluorescence and label free nanoscale imaging permit quantitative 3D spatial temporal detail of individual cellular components that will define the next generation of nanoscopy to study cell migration.

In this paper, we demonstrate for the first time, three imaging modalities (ROCS^16, 17^, HiLo^23, 24^) combined in an oblique illumination system using a single laser source, a diaphragm and 2 cameras. While ROCS and HiLo are constructed on an oblique scanning system, quantitative interferometric imaging has not been reported to use an oblique illumination system before ^25^. Furthermore, BF and DF ROCS improve imaging contrast of scatterers but do not produce visible interference intensity fringes that is present in interferometric imaging such as iSCAT ^15-17, 26^ or reflection interference contrast microscopy (RICM) ^27-31^. Images from these interferometric techniques display characteristic varying interference fringes produced from scattered and reflecting light from the sample and glass surface. Here, we report a new interferometric imaging that is achieved using the ROCS platform. At 22° oblique angle for a rotating beam, we were able to retrieve distinctive intensity fringes ^32^ which can be used to quantify axial position of scatterers (nanoparticles, organelle in living cells). We term this imaging mode as interferometric ROCS. In the following sections, we first outline the optical design (scanning angle and Fourier amplitude filter) necessary to obtain interferometric, BF and DF ROCS imaging in an oblique illumination system. Secondly, we carried out calibration experiments to measure and compare the imaging sensitivity of interferometric, scattering and fluorescence signal using 20 nm, 100 nm and 200 nm nanoparticles along axial and lateral direction ^19^. We then applied the combined imaging platform we study two cell types that actively migrate in response to vessel injury. Thirdly, we establish biological goals for interferometric, scattering and fluorescence imaging experiments to achieve. In the first imaging experiment, we examine platelets that are thought to generate filipodium protrusions that spans several platelet lengths (∼ 5-10 µm) ^33, 34, 35^ to form a biomechanical plug ^36, 37^. Platelet can direct their migration to scavenge adhesive molecules bound to a stiff substrate ^4^, that links their role as cellular scavengers to collect molecular agents. In the second imaging experiments, we carried out imaging on endothelial cells that migrate to repair the inner lining of damaged vasculature ^38^. We used fluorescence imaging to study real-time platelet migration on glass surfaces coated with fluorophore-labelled fibrinogen and crossed-linked fibrinogen-fibrin, in the presence and absence of Bovine serum albumin (BSA)-common protein in blood. Mitochondrial shape and signaling in endothelial cells are known to influence cell migration, proliferation, and tissue reorganization ^39-41^. The mitochondria 3D network needs to fragment (*i.e*. by mitochondria fission) ^41, 42^ to smaller units that can be more readily transported across the cell ^40^ to provide energy for cell movement. Using fluorescence labelling of the mitochondria, we identify and measured the axial motion of mitochondria in migrating endothelial cells. We show that regions of the mitochondria between the leading edge and nucleus exhibit higher axial motion. In the two biological imaging experiments, we focused on correlating the spatial organization of membrane, organelles in subcellular regions of cells ^39, 41, 43^ along edge of cell membrane (leading and rear edge) ^44^ that orchestrates cell migration. Combined oblique imaging with interference and scattering permits us to quantify membrane fluctuations of migrating cells platelets and endothelial cells at the leading and rear edge and leverage fluorescence to identifying surface molecules or intracellular organelle that regulate cell migration.

## Results

### Oblique angle and amplitude Fourier filter for interference and scattering imaging

Oblique illuminated azimuthal scanning microscope systems are frequently used in ROCS (BF and DF) ^16, 26^, HILO ^23, 24^ and COSI ^45^ to achieve high contrast imaging for both fluorescence and scattered light but does not achieve quantitative depth imaging using interferometry. In ROCS ^26^, samples are typically illuminated with a rotating beam at incoming oblique angles greater than 60°. At this angle, ROCS do not exhibit interference intensity fringes present in interferometric imaging ^19^, that are used to detect nanoparticles e.g., gold, purified protein, virus particles ^27, 28, 46, 47, 48^. We empirically observed that the oblique scanning angle and the position of an amplitude filter (diaphragm) placed at the imaging back focal plane (BFP) is critical to toggle between interferometric, BF and DF ROCS. The configuration of oblique angles and diaphragm determines the signal to noise ratio (SNR). Here, we refer to these ROCS imaging modes as “scattering” and “interferometric” imaging, respectively. In quantitative interferometric imaging (e.g. iSCAT, RICM) ^27-30^, the interference signal is primarily optimized by balancing intensities reflected from a glass coverslip with weak scattered light from nanoparticles or the biological sample using apodised optics ^25, 29^. To visualize quantitative interference signal, the intensity of reflected and scattered must be tuned to obtain the optimal contrast of interference fringes. The optimal imaging contrast in interferometry-based methods is achieved by modifying the incidence angle and diameter of the amplitude filter. In supplementary Figure S3, we showed that the amount of back-reflection increases exponentially at larger angles (> 40°, Supplementary Figure S3), with a shift of oblique angle from 20° to 60°. Unlike traditional iSCAT/RICM that retrieves interferometric intensity fringes at normal incidence, interferometric ROCS retrieves interference fringes at lower oblique angles of 22°. To effectively switch between interferometric, scattering and fluorescence microscopy, we need to simply switch between the scanning angles as shown in Figure. 1A, where we projected 3 annular scanning illumination paths (red, blue and green) implemented on a single high numerical aperture (NA) objective lens of 1.49. The interference of back-reflection and the scattered light is collected here is at an illumination angle of 20°-35° (∼N.A. = 1.49). This differs from ROCS where the authors retrieved the interference signal at a much larger angle (N.A. = 1.2 and 1.46).

**Figure 1.**
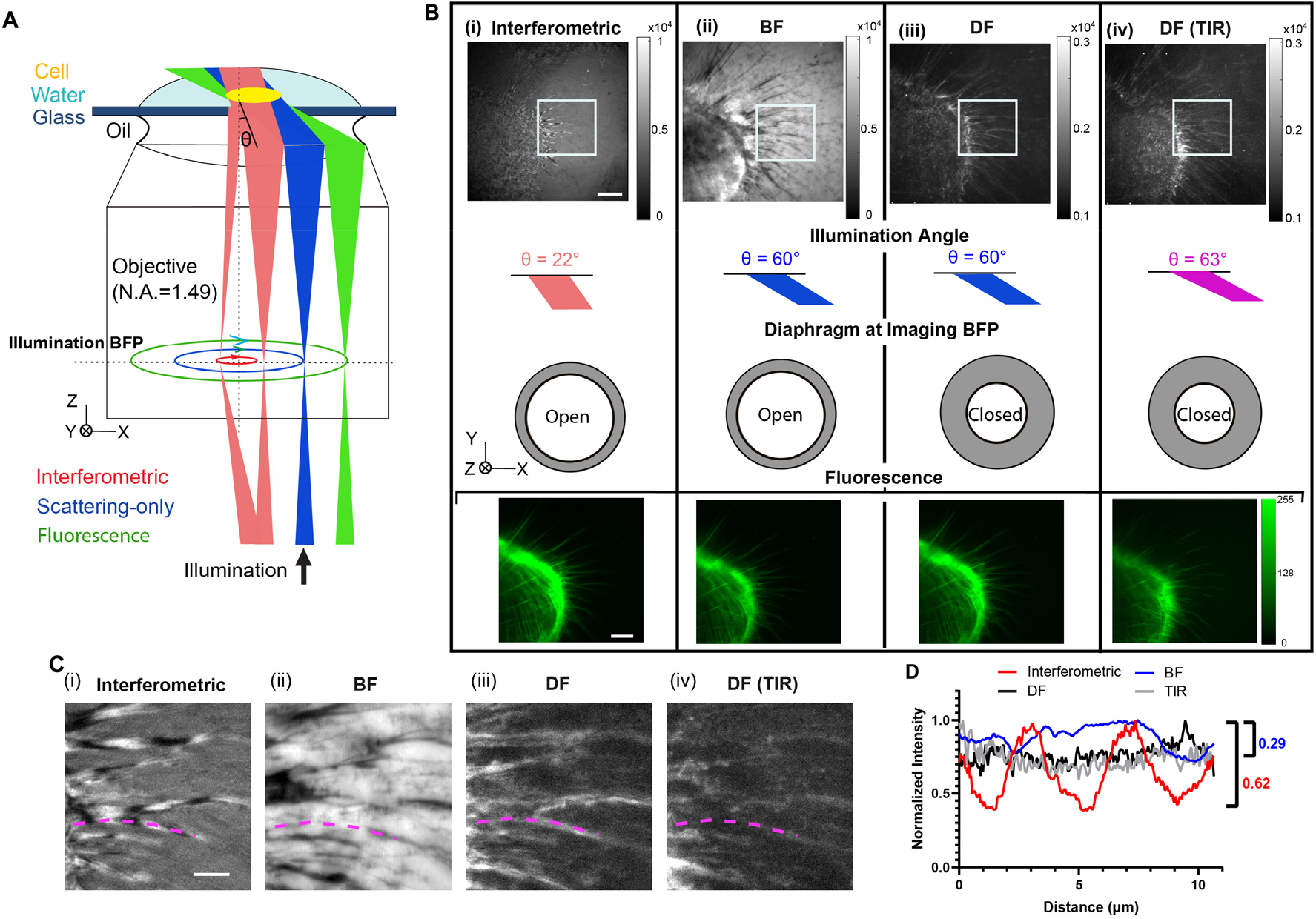
Combined Interferometric, Scattering and Fluorescence Imaging. (**A**) An oblique beam (with angle θ corresponding to optical axis) is radically scanning at back focal plane (BFP) of the 1.49 NA objective. When θ = 10° to 40° (red), the interference of back-reflection from the coverslip and scattered light will be collected by the detector, and when θ > 50° (green), the back-reflection will be blocked by a diaphragm at the imaging BFP that only allow pure scattering to be detected (darkfield). The fluorescence signal (blue) will be collected simultaneously at corresponding angles without blocking due to the dual camera detection scheme. (**B**) Label-free and fluorescent imaging using interferometric, BF, DF and TIR ROCS imaging of HMVEC cells (labelled with Alexa Fluor 488 conjugated Phalloidin). Scale bar = 5 μm. The illumination angles (22°, 60° (HiLo) and 63° (TIRF)) and diaphragm opening at the imaging BFP are illustrated for each modality. (**C**) Zoomed views of the highlighted filopodia region in (**B, i-iv**) for each modality and (**D**) plot of the fringe intensity profile along a filopodium indicated in D (i-iv) (magenta dashed line). Scale bar = 2 μm.

Supplementary Figure. S1 describes the optical design and setup in detail. During imaging, a single laser beam (wavelength, λ= 488 nm) is directed onto a two-axis galvanometer and conjugated onto the back focal plane of the objective lens. The two-axis galvanometer allows precise adjustment to different scanning radii which in turn selects the imaging mode (Scattering, Interferometric, and Fluorescence). While fluorescence imaging (green) can be obtained from any azimuthal scanning pattern ^24, 49^, interferometric and scattering-only images ^16, 29^ are separated by two scanning radii (red, blue). Experimentally, this can be determined and preset, as shown in Fig. 1A on conjugated back focal planes (BFP). Here, interference contrast is maximized without using specialized partially reflective mirror ^*29*^. As shown in Fig. 1B, at higher azimuthal oblique angles (>50°), one can choose between brightfield and darkfield ROCS by opening (BF) or closing (DF) the diaphragm placed in the BFP to remove reflected intensities (*i.e*. from coverglass and particles). Using the oblique illumination angle alone, we can achieve interferometric imaging at an oblique angle θ of 22° and scattering mode at an illumination angle greater than 50° (Figure 1B, S2, S3, S4). The iris diaphragm has zero transmittance on the edge and will not induce phase shift. We then tested the combined imaging of fixed HMVEC cells adhered on a glass coverslip. Fig. 1B (i) to (iv) shows a sequence of interferometric, scattering (BF and DF ROCS) and fluorescence images of F-actin (phalloidin)-labelled HMVEC cells under different oblique illumination angles from 22° to 63°. While changing from 55° to 60°, we observed the inversion of scattering and background signal (Fig. 1B (i, ii)) from brightfield to darkfield. Fluorescence images taken at oblique illumination angles (θ = 60°, 63°) displayed increased contrast than at θ = 22°. Fig. 1C shows the magnified view of filopodia indicated in Fig. 1B and Fig. 1D, the measured intensities along the marked filopodium protrusion. Restricting the diaphragm diameter leads to contrast inversion of filopodia from BF to DF ROCS (θ = 60°). In BF and DF ROCS, interference intensity fringe are no longer observed. In interferometric ROCS, interference intensity fringes are distributed along a filopodia (Fig. 1D, Movie 1). We then quantify the intensity fringe against the intensity signal from BF and DF ROCS is shown in (Fig. 1D) after normalizing the intensities. This comparison informs us that the intensity variations of interference fringes from interferometric ROCS is distinctively different from that of BF and DF ROCS.

Interferometric scattering signals, shown in Fig. 1C i) and D) provided a negative contrast ^30, 50^ because of the cosine term in the interferometric component 2*E*_*r*_*E*_*s*_*cos*ϕ^51^, where *E*_*r*_, is the reflected electric field from the coverslip, *E*_*s*_ is scattering electric field from the sample and ϕ is the relative phase difference between *E*_*r*_ and *E*_*s*_. Scattering cross-section of the sample (*E*_*s*_) will scale nonlinearly with particle size, where *E*_*r*_ will scale with just reflecting angle. In other words, a smaller object will thereby have lower scattering (*E*_*s*_) and *E*_*r*_ changes with just angle. As with any interference signal, the balance of *E*_*r*_ and *E*_*s*_ are necessary for high fringe contrast. Because of sample variability, the visibility of the interference fringes in an interferometric scattering signal require the balancing between *E*_*r*_ and *E*_*s*_ as shown in Fig. 2.

**Figure 2.**
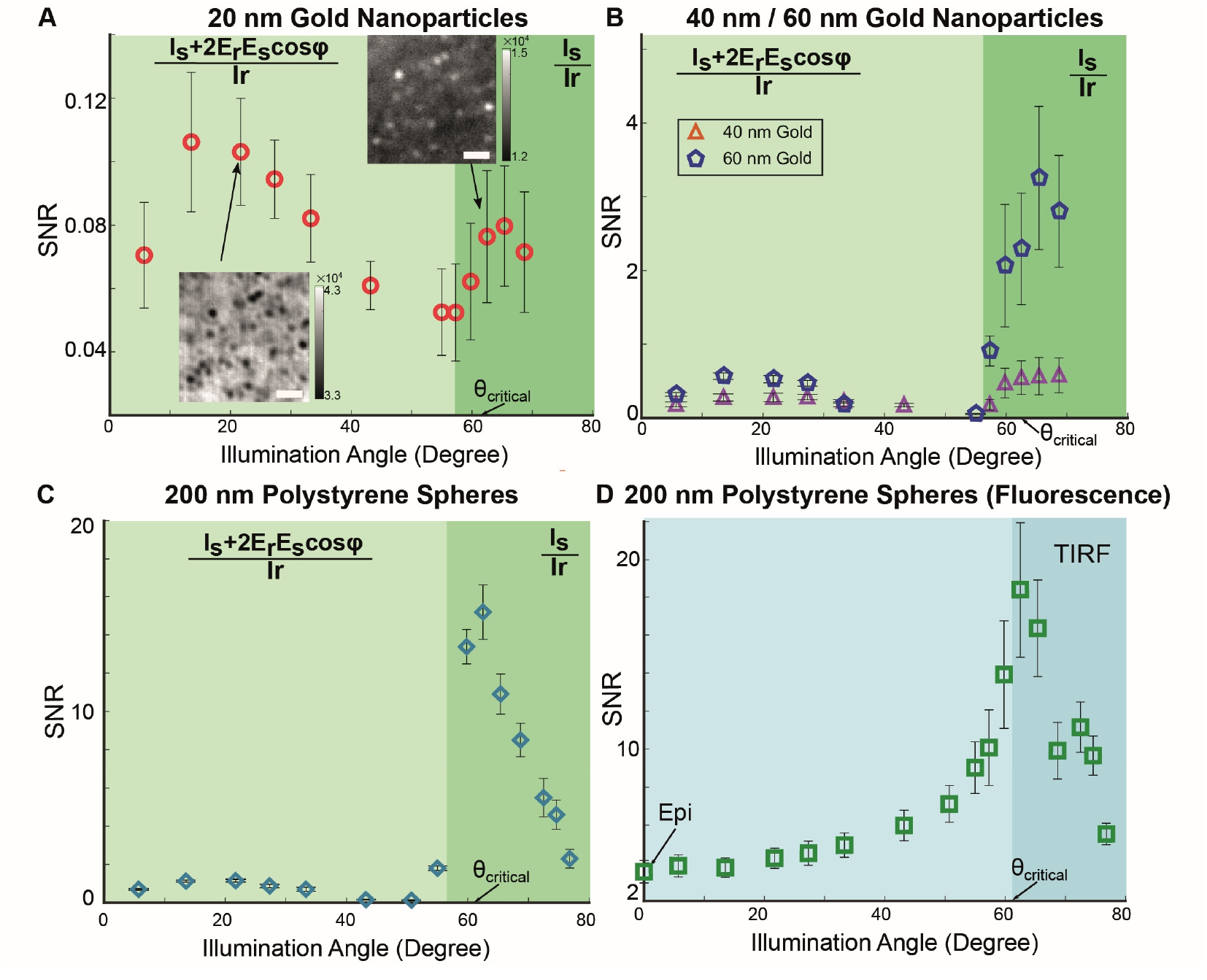
Quantification of signal to noise ratio of nanoparticles ranging from 20 – 200nm (diameter) with scattering and fluorescence detection. SNR of (A) 20 nm, (B) 40 nm and 60 nm gold nanoparticles and (C) 200 nm polystyrene using scattering detection under different illumination angle ranging from 5° to 70°, where diaphragm is set to block back-reflection at 57° and the critical angle is at 63°. Inlet images in (A) are interferometric and pure scattering signal of the particles at 22° and 63° respectively. Scale bar = 1 μm. (D) SNR of the 200 nm polystyrene with fluorescence detection, where the critical angle is at 63°. Error bar = standard derivation from measurement of at least 50 nanoparticles.

We next investigated detection limit of each imaging mode, interferometric, scattering and fluorescence using gold and polymer nanoparticles. Fig. 2A to D compares imaging contrasts of interferometry, scattering and fluorescence using signal to noise ratio (SNR) by scanning oblique angle from 5.7° to 76.8 °. Details of SNR for both negative and positive contrast is detailed under Method section. We chose a range of gold nanoparticles with mean diameter 20± 6.3 nm, 40± 8.9 nm and 60± 17 nm and fluorescence beads of 200 nm (as specified by the manufacturer). The nanoparticle diameters were chosen because cross-section of gold scales exponentially below 100 nm, modeled in Fig. S3, which influence the scattering intensity (I_s_). Fig. 2A and 2B shows gold nanoparticles with 20nm, 40nm and 60 nm under various oblique illumination angle. For gold nanoparticles, we only compare the interference and scattering signal in water. SNR results for 20 nm gold with a low scattering cross-section of 1.32 nm^2^ (∼1.6% of total surface area) is much higher for interferometric than scattering imaging. Fig. 2A show that scattered SNR (Is) nanoparticles exhibit an SNR of 0.08 ± 0.022 whereas the interferometric 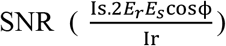^51^ is about 37% higher (0.11 ± 0.022) Conversely, we showed that an increase of the scattering cross-section by orders of magnitudes, ∼ 81.3 nm^2^ (40 nm gold nanoparticle) and 791 nm^2^ (60 nm gold nanoparticle) results in higher scattering SNR when compared with interferometric SNR as shown in Fig. 2B. Fig. 2B showed that scattering SNR reached a value of 3.25 ± 0.97 to 0.566 ± 0.05 which are higher than the value of interferometric SNR that hovers around 0.278 ± 0.05 to 0.60 ± 0.27. Our finding shows that interferometric SNR (2*E*_*r*_*E*_*s*_*cos*ϕ) ^51^ can play a significant role to improve imaging contrast when particles have small scattering cross-section of around 1.32 nm^2^. Next we compare particles size with diameter of 200 nm. Fig. 2C and D compares the interferometric, scattering and fluorescence SNR of 200 nm polystyrene beads. Fig.2C and D compare the interferometric, scattering and fluorescence SNR of 200 nm polystyrene beads with a scattering cross-section of 4866 nm^2^ that is over 3 orders of magnitude higher than observed for 20 nm gold nanoparticles. Scattering SNR in 200 nm polystyrene beads has value of 15.19 ± 1.42 that is close to 10-fold higher than interferometric SNR with value of 1.13 ± 0.05. To briefly summaries, the diaphragm placed at imaging BFP acts as an intensity apodizer. This meant that at oblique illumination angles from θ=10° to 40°, we achieve interferometric imaging. Beyond oblique angles of θ>50°, scattering imaging is achieved. By simply shifting the oblique illumination angle, nanoparticles of sizes from 20 nm to 200 nm can be readily imaged by interferometry. Azimuthal scanning removes coherent speckle noise and artifacts that are present in interferometric imaging, scattering and fluorescence ^16, 49^. In addition to the illumination angle, the polarization of illumination and sample-scattered light can also affect the interferometric signal.

Interferometric SNR would naturally possess axial information that can be used to retrieve a 3D point spread function ^46^. This permits interferometric signals to track gold nanoparticle axially ^28, 46^ that can be developed for tracking cell membrane protrusions ^52^. To calibrate this, we investigated axial interference SNR using the same nanoparticles sizes ranging from 20 nm, 60 nm and 100 nm gold as well as 100 nm polystyrene. All the particles are fixed on a thin coverglass which is then placed on a piezo nanostage (P-736, Physik Instrument) as shown in Fig. 3A. The sample illuminated obliquely at 22° is swept axially through the objective focus at an interval of 10 nm step sizes. As shown in Fig. 3B and C, for 60 nm and 100 nm gold nanoparticles (higher scattering cross section of 7620 nm^2^ and 81.3 nm^2^), the axial interferometric intensity displays a characteristic sinusoidal intensity gradient. On the other hand, when scattering cross section is reduced by an order of magnitude to ∼1.32 nm^2^, the intensity gradient followed a parabolic curve as presented Fig. 3D. The parabolic curve in 20 nm gold nanoparticles indicate that destructive interference (*cos*ϕ) ^51^ is responsible for high axial resolution for nanometer gold scatterers. Axial interferometric SNR for membrane tracking used the linear region calibrated with 100 nm polystyrene sphere that has a refractive index close to actin (∼1.57) ^53^. Using the calibration of the interferometric signal (Figure 3), we estimate the axial resolution with the equation: height difference/ΔI. The axial resolution therefore scales to the sample scattering cross section. A smaller scattering cross section yields lower axial resolution (0.3 nm for 20 nm gold beads) and higher resolution with larger scattering cross sections (0.05 nm for 100 nm gold beads). In summary, Fig. 3B to E showed that the total axial tracking distance of interference for all particle sizes ranges between 0.67 to 0.7 µm.

**Figure 3.**
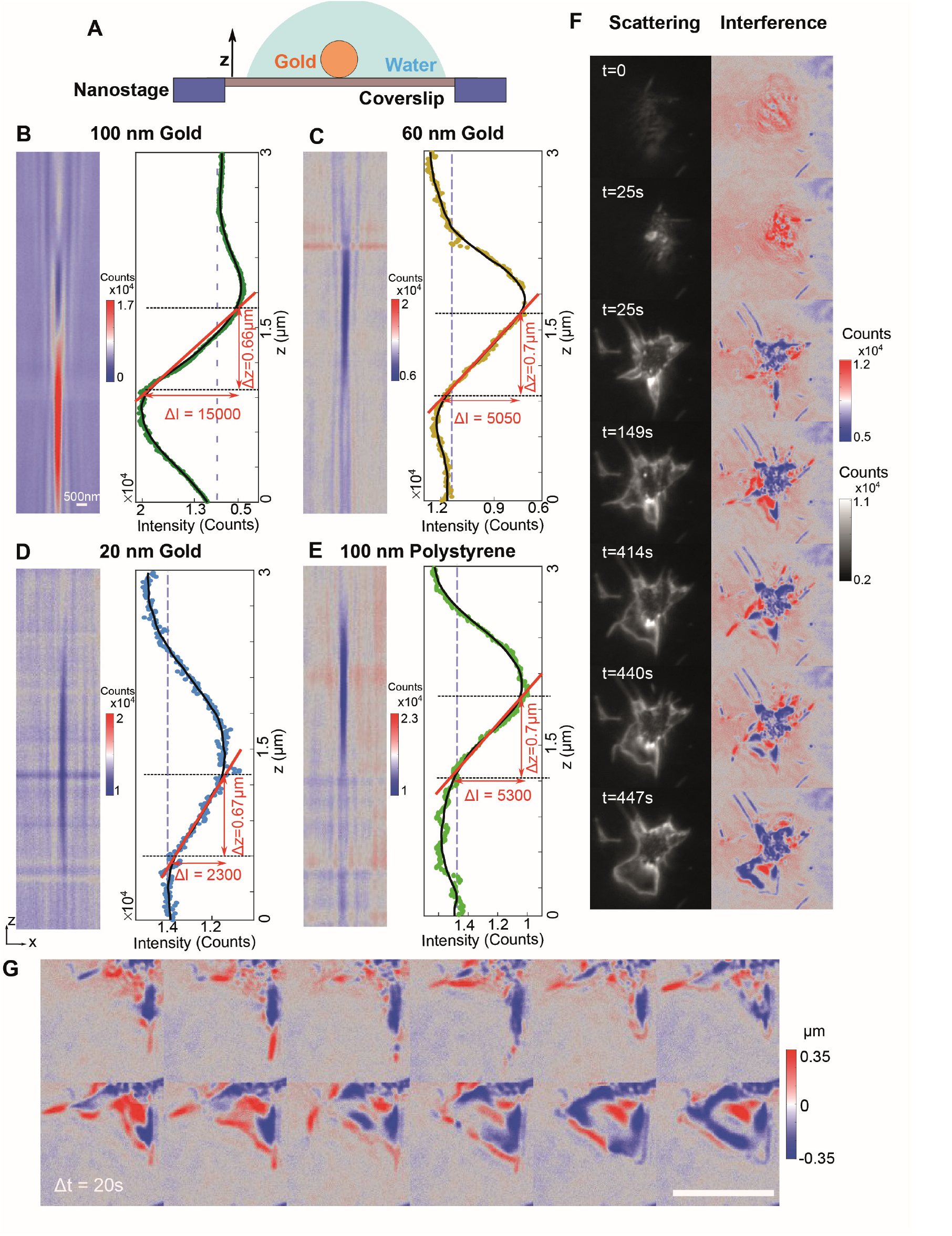
Calibrating axial intensity profile and displacement with nanoparticles from 20 nm to 100 nm. (A) To record the intensity profile of gold/polystyrene nanoparticles, we firstly dried the particles on a coverslip and then immersed with water. The coverslip was place on a piezo nanostage that allows axial sweeping of the sample across the focus at 10 nm steps. The axial intensity map along with the line profile of gold nanoparticles at (B) 100 nm, (C) 60nm and (D) 20nm, and (E) polystyrene nanoparticles at 100 nm are plotted as scattered plot, the black line indicates the moving average of 30 points. The red line represents the relation between intensity (ΔI) and axial displacement (Δz). (F) Simultaneous recording of both scattering and interferometry over a period of 447 seconds showing early platelet adhesion, filopodia and lamellipodia formation and spreading on fibrin surfaces. (Movie 2) Scale bar = 5 μm. (G) Axial displacements of the membrane protrusions at 20 seconds time intervals. (Movie 3)

### Imaging of single platelet migrating at the nanoscale

Recent live imaging studies of platelets have showed different biomechanical actions associated with platelet membrane protrusions. Platelet filopodium was observed to make “hand over hand” actions on fibrin fibers^54^. Platelet have been seen to migrate with formed lamellipodium when exposure to pathogens on coated fibrinogen surfaces. We previously demonstrated that high speed label-free platelet imaging, with rotating coherent scattering (ROCS) ^16, 55^, captured unlabeled platelet generating rodlike-extension filopodia instead of thin-sheet lamellipodia to directly attach onto single collagen fibril under arterial fluid shear ^45, 56^. ROCS alone is insufficient to investigate other platelet morphodynamics along the axial direction as well as membrane surface signaling processes. We now overcome the limitation by using combined fluorescence, interferometric and scattering imaging to study platelet lateral and axial membrane protrusions dynamics on glass surfaces coated with fibrinogen or fibrin (cross-linked fibrinogen).

For each coated surfaces, we added either add or omitted BSA to study platelet migration. Details of sample preparation and image analysis are available in supplementary materials-Methods and sample preparation and Figure. S5. We first evaluate simultaneous scattering and interferometric imaging on platelet arriving onto fibrin coated surface. Unlike traditional RIC imaging, we can capture platelet adhesion from initial contact to initial stages of spreading and adhesion on fibrin-coated glass surfaces as shown in Fig. 3F. Fig. 3F shows data of simultaneous recording of both scattering and interferometry over a period of 447 seconds (see Movie 2). While interference signals that only occurs close to the coverslip, the scattering signal suppressed background to capture floating platelet prior to adhesion at high contrast (without postprocessing). The scattering signal is sufficiently high for automated segmentation and tracking. The interferometric signal is used to quantify axial displacements on those membrane protrusions as shown in Fig. 3G at 5 seconds time intervals. Fig. 3G tracks the first instances of filopodia formation during adhesion before a lamellipodium is formed (see Movie 3). To visualized migrating tracks with and without BSA, we labelled fibrinogen and fibrin with fluorescence label (Alexa Fluor 488) using previous published protocol ^4, 57^. Here, migrating tracks are shown by a lost in fluorescence due to uptake of fibrin or fibrinogen coating on glass surfaces. The tracks are imaged using fluorescence imaging.

Fig. 4A(i) to (iv) shows interferometry, scattering and fluorescence, using BSA treated sample <u>(see Movie 4)</u>. Adherent platelets are seen to internalized fibrinogen and fibrin. A bright fluorescence spot can be seen located intracellularly in the platelet using fluorescence. Using scattering and interferometric imaging, platelets’ cellular morphology in 3D can be quantified using standard cell segmentation and profiling tools in open source program *Cellprofiler* ^58^. We sorted the imaged platelets (n_fibrin_=427 and n_fibrinogen_ 510) against two different classifiers namely area (µm^2^) and solidarity. Area is meant to show cell spreading. For platelets with a larger spread could indicating flat membrane protrusion resembling lamellipodium. Solidarity is defined by irregularity along the cell boundaries. Platelets with highly number of filopodium protrusion (finger-like) would score low on the solidarity point. Additional information on cell segmentation and morphological profiling is detailed in Fig. S5. Fig. 4 B (i) show the classifiers (area and solidarity) and the percentage of migrating platelets on fibrin and fibrinogen coated surfaces. The results showed that without the inclusion of BSA, no migration is observed (no visible tracked due to loss of fluorescence). Overall, Fig. 4B (i) showed that there are around 4 folds more platelets migrating under fibrin coated. Conversely, platelets would generally migrate on fibrinogen coated surfaces. Using the classifiers, we can further separate the morphological profiles of the migrating and non-migrating platelets as shown in Fig. 4 B ii) and iii). Platelets on fibrin coated surface showed comparatively smaller surface area than fibrinogen coated surfaces. On the other hand, on the solidity scoring, there is no significant variation amongst the platelets either migrating or non-migrating. Overall, this meant that the platelet adhesion under spreading conditions would generate highly heterogenous morphology profiles. Such heterogenous membrane protrusion differs from our previous observation of platelet under flow on collagen matrix where scattering imaging only displayed filopodia formation and could have important implication of thrombus-based fibrinolysis. With scattering and interferometric signal, we carried out automated axial tracking of lamellipodium of a migrating platelet on fibrin coated surface as shown in Fig. 4C (i) (Movie 5, 6) that indicated a fluctuation from peak to peak height displacement of around 0.13 µm as shown in Fig. 4C (ii). In Fig. 4C (iii), we then compared both the leading and rear edge of the migrating platelet by plotting the overall axial fluctuations. The leading edge showed a normal distribution of membrane displacement entered at 0 (*i.e*. no protrusion or retraction). However, the rear edge exhibited unequal membrane fluctuations of -0.13 μm (retraction) to 0.13 μm (extension), shown as two peaks in Figure 4C, iii). The heterogeneity of membrane fluctuations may reflect unequal membrane tension during cell migration. Out of phase fluctuations indicate asynchronous behavior of cell migration inferring differences in adhesion strength ^59^ between the leading and rear edge.

**Figure 4.**
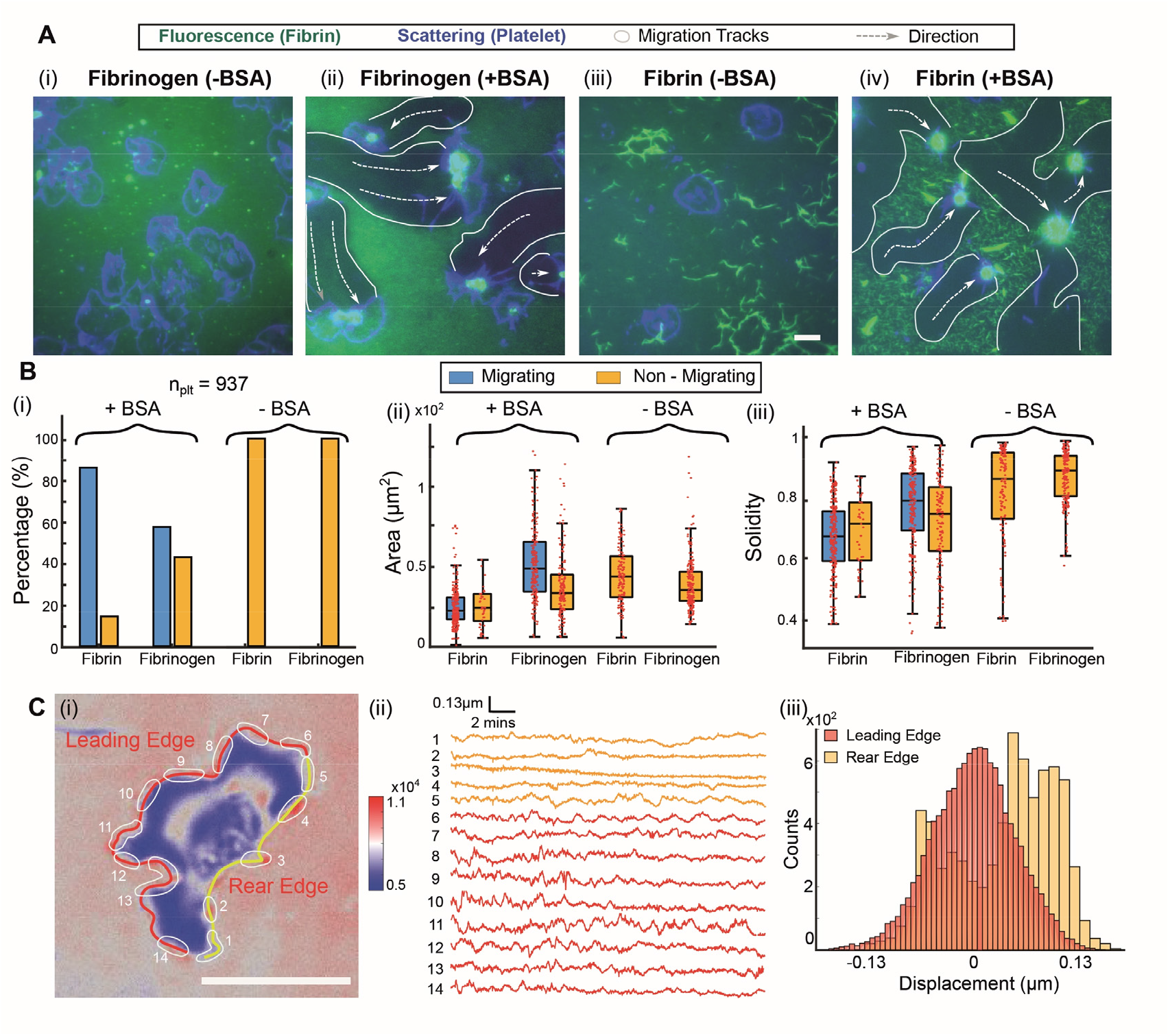
Tracking platelet migration on fibrin/fibrinogen coated surface. (A) Combined fluorescence and scattering signal of single platelet migration on fibrin (Movie 4) and fibrinogen coated surfaces with/without BSA. Green channel and blue channel are fluorescence and scattering signal respectively. The trajectory of platelet movement is highlighted with white broader with white arrow indicating predicted direction of migration. (B) Quantification of platelet migration and morphology (area and solidity) with four types of coating. Platelet morphology is quantified through cell profiler. Number of platelets quantified: n = 427 and n = 510 for fibrin and fibrinogen coated surface respectively. (C) (i) Interferometric signal of a migrating platelet on fibrin coated surface. (ii) The axial displacement along the leading edge and rear edge is tracked during 25 minutes of spreading, with orange and red represent rear and leading edge respectively (Movie 5, 6). (iii) Histogram showing the counts on axial displacement of rear and leading edge.

### Imaging of Nanoscale dynamics of Mitochondria in migrating cells

The mitochondria form a 1 micron-thick tubular network and trafficked along the cytoskeleton to distribute energy production for cell motility ^39, 60^. Visualizing this mitochondrial organization in real-time relies on fluorescence labelling, which is prone to photobleaching and does not resolve the seconds-timescale remodeling of the mitochondrial 3D network necessary for cell migration ^61^. Mitochondria motion is therefore challenged by the 3D space they are in: thicker cell space near the nucleus, but interlinked with the endoplasmic reticulum network ^60^ or thinner cell space at the leading edge (*i.e*. lamellipodia) of a migrating cell. Considering scattering and interferometric imaging provides both lateral and axial (Figure 3) nanometer imaging resolution, we explored whether both imaging modalities resolve mitochondria 3D structure and displacement in migrating cells. For this, we employed microvascular endothelial cell line labelled with Mitotracker Green plated on glass-bottom culture dishes, which are commonly studied in scratch wound assays to investigate migration during vascular injury ^38^. Our initial results found significant photobleaching of Mitotracker fluorescence resulting in 30% of fluorescence intensity over 3 minutes (Supplementary Figure. S6). To overcome this, we synchronized the laser emission to the scanning frequency to generate a pulse of 0, 2, 10 or 20 Hz (Figure 5A and supplementary methods). We found that increasing the laser pulse rate to 2 Hz reduced photobleaching by 15% and maximum reduction of photobleaching by 25% with 10 and 20 Hz pulse (Supplementary Figure. S6). Hence, to minimize photobleaching and phototoxicity we conducted all mitochondria imaging experiments with 20 Hz laser pulsing.

**Figure 5.**
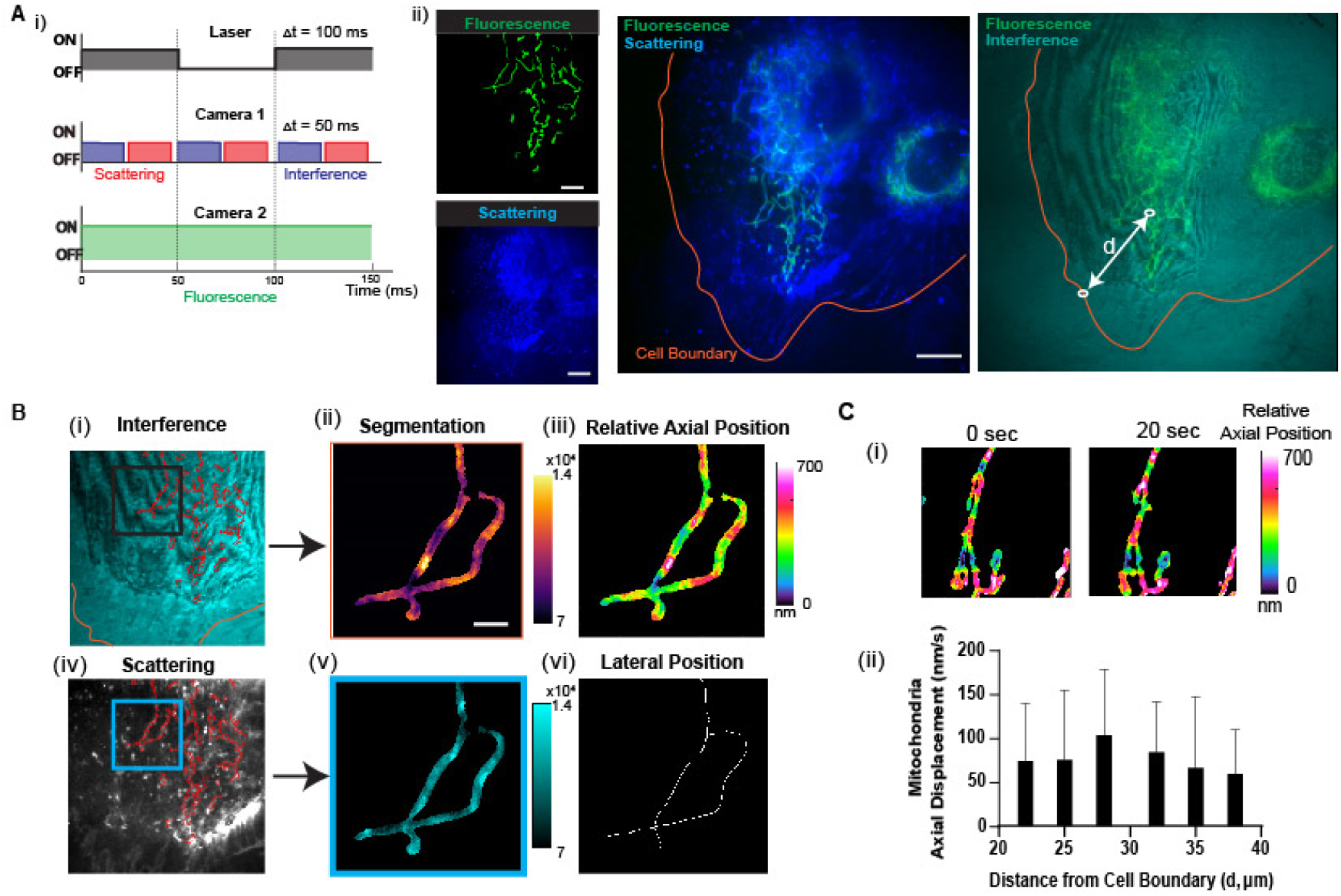
Tracking of mitochondria axial position in endothelial cells. **(A, i)** Diagram showing laser pulse synchronized to camera 1 (scattering and interferometric imaging) and camera 2 (fluorescence imaging) to reduce photobleaching. (ii) The fluorescence, scattering and interferometric images of HMVEC cell labelling of mitochondria with Mitotracker Green. (B) The mitotracker fluorescence signal for (ii, v) masking and segmenting mitochondria in the (i-iii) interferometric and (iv-vii) scattering images. (iii) The interferometric intensity is converted to relative axial position using the calibrated axial intensity profile from Figure 3 and (vi) scattering image provides lateral position of the mitochondria. (C, i) The relative axial position of mitochondria in HMVEC cells retrieved from interferometric signal and (ii) plot of axial mitochondria displacement as a function of distance from the cell’s leading edge. Data are S.D. of n=9 data points.

Under scattering and interferometric imaging, we observed tubular structures extending 25 µm outward from the perinuclear region (Figure 5A (ii)). These structures split and converged along its length, consistent with a mitochondrial network under persistent fusion and fission. Interferometric imaging revealed fringe shifts that were unique to the tubular structures and not the entire cell. To confirm these were mitochondrial networks, we stained HMVEC cells with a mitochondria-specific fluorescent dye, Mitotracker Green (Figure 5A (ii)), which confirmed the observed tubular structures were mitochondria. Figure 5B (i) and (iv) shows that the Mitotracker fluorescence signal is used as a mask to segment mitochondria in the interferometric (Figure 5B (ii)) or scattering image (Figure 5B (v)). Figure 5B (iii) shows a relative axial position map of individual mitochondria filaments retrieved calibrating the fringe shift to axial distance in Figure 3. The scattering image complements the axial information, providing the lateral position of the mitochondria (Figure 5B (vi)).

Our quantitation of mitochondria relative axial position reveals a morphology with multiple bends across its length. We next measured the variation of mitochondria axial movement across a migrating cell (Figure 5C and see Movie 7). Figure 5C (i) shows the axial shifts across a mitochondrion of 45 µm length taken 20 seconds apart. Here, axial shifts are determined by the intensity fringes relating to the periodic phase shift, producing a relative height measurement of structures. We observed axial shifts of up to 350 nm within a length of 2 µm and regions of increased axial motion that was centered in the middle (∼20 µm) of the mitochondrion. Interferometric imaging also revealed the mitochondrion stretched up to 15 µm away from the leading edge of the cell, where the cell actively formed membrane protrusions and expanded its boundary by 3.8 µm (see Movie 7). Having found a variance in mitochondria axial motion, we examined if proximity to the cell’s boundary affected mitochondrial motion. Preliminary evidence suggests that mitochondria bind to focal adhesions ^62^, which may constrict mitochondria motion. Figure 5C (ii) shows the axial displacement of the mitochondria in Figure 5C (i), measured between 22 – 38 µm distance from the leading edge of the cell. Mitochondria axial motion ranged between 10 nm/s to 100 nm/s and was maximal at 28 µm away from the cell boundary. These results support our observation from Figure 5C (i). We suggest that mitochondria tethered to the endoplasmic reticulum network and focal adhesions may play a role in regulating mitochondria motility ^39, 40^. Through combined interferometric, scattering and fluorescence imaging of the mitochondria and cell membrane, we additionally visualized the distribution of mitochondria centered around the nucleus of the cell and its entry into the nucleus (Movie 8).

## Conclusion

Nanoscale live cell imaging illuminates our fundamental understanding of cell migration. Existing nanoscopic imaging tools are broadly classified into separate fluorescence ^14^, scattering ^16, 55^ and interferometric platforms.^19, 20^. We demonstrated a direct means to combine scattering, interferometric and fluorescence imaging modalities system using a single laser source with azimuthal laser scanning and dual digital cameras. Most importantly, we identified that a Fourier amplitude filter is necessary to perform interferometric imaging ^49^ using azimuthal scanning that is able to balance intensities reflected from a glass coverslip with weak scattered light from nanoparticles or biological sample without using apodised optics ^25,29^. In ROCS, the aperture of the diaphragm needs to be changed when switching imaging mode whereas the aperture of the diaphragm here is fixed. Hence, the combined ROCS imaging platform can switch between interferometric and scattering imaging modes for simultaneous multimodal imaging. In the current combined imaging system, we do not use background subtraction that is used in ROCS. We are able to obtain high contrast imaging (sensitivity up to 20 nm gold nanoparticle) that does not need background removal in real time. The combined oblique illumination modality unifies interference, scattering and fluorescence ROCS into a single unit and has shown to be applicable for a multitude of imaging experiment that investigate spatial organization of intracellular organelles ^63^, ligand or lipid coating on substrates surfaces ^64^ that are necessary for cells to migrate ^39^. Interferometric signals possess an axial resolution of lower than 100 nm.

The next step is to apply the combined imaging platform to study membrane receptor clustering ^65^, intracellular calcium flux ^66^, actin polymerization ^67^ while continuous tracking morphological changes in cell migration assays. We further anticipate that the combined imaging can possibly resolve nanoscale membrane curvatures ^68^, membrane pores^69^, transmembrane proteins and receptors,^70, 71^ as well as external mechanical forces^72^ that influence the formation of membrane protrusions (lamellipodia, ruffles, filopodia). While we have shown that the combined imaging technical can be applicable to both anucleate and nucleated cells, the limitation of interferometric signals when combined is the limited imaging depth of ∼ 0.7 µm. Finally, the current interferometric imaging do not use any form of post-processing and we postulate that detection sensitivity could be further enhanced by using ratiometric subtraction.

## Acknowledgement

We thank Ms. Sarah Hicks and Mr. Junxiang Zhang for providing washed platelets and fibroblast samples.

## Contribution

W.M.L. conceived and supervised the project. Y.Z. built the imaging system and performed all platelet experiments. Y.J. L develop live cell imaging chamber and conducted intracellular imaging on endothelial cells, H. L developed the automated image tracking method, T Xu advice on optical design, Y.L.T. prepared washed platelet samples, E.E.G advised on platelet experiments, V D-A and C Parish advised on endothelial cell experiments, and. Y.Z. carried out the data analysis along with H.L (*Intensity tracking*) and C.L. (*Cell profiler*). Y. Z, Y.J.L and W.M.L. wrote the manuscript with input from all authors.

