## Supplementary Files for "Combined Scattering, Interferometric and Fluorescence Oblique Illumination for Live Cell Nanoscale Imaging"

Address for correspondence

Dr W M Lee

The Australian National University,

Canberra ACT 2601, Australia

*

### Methods and sample preparation

##### A. Dual camera detection for a single azimuthal scanning- oblique illuminated beam

The imaging system is constructed on an inverted microscopy platform (RAMM, ASI) using a linearly polarized 488 nm laser (Stradus 488-150), as shown in Fig. S1. A two-axis galvo mirror (GVS212/M, Thorlabs) is conjugated to the imaging plane of the system so that the laser beam is focused on the back focal plane of the objective lens (Olympus, 60X, NA 1.49, oil immersion) to allow wide-field illumination. The fluorescence and scattering signal is separated using a dichroic mirror (FF505-SDi01-25x36, Semrock) and collected by a CCD (Retiga 4000R, QImaging) and a sCMOS (PCO edge 4.2), respectively. To achieve oblique illumination and azimuthal averaging, a galvo mirror (GVS212/M, Thorlabs) was used to radically scan around the optical axis at the back focal plane of the objective. We also synchronized the capture rate of the sCMOS and the scanning frequency with a data acquisition card (PCIe-6374, National Instruments) so that one complete cycle of an azimuthal scan can be captured and averaged simultaneously. The laser on/off trigger was also synchronized with the scanning frequency to create a pulsed illumination and reduce photobleaching (see Fig. S5).

The switching between interference signal and pure scattering detection is achieved through tailoring the oblique illumination angle by controlling the scan radius at the BFP of the objective and Fourier amplitude filtering by placing a diaphragm (Thorlabs, ID15/M) at the Fourier plane (conjugated to the back focal plane of the objective) as shown in Fig. S1. The diaphragm modifies the BFP aperture and the detection N.A. of the scattering detection (N.A. = 1.27, illumination angle = 57°) so that the back reflection can be blocked to collect pure scattering signal. The detection N.A. of the fluorescence signal is not limited by the diaphragm and will always be equal to the N.A. of the objective (1.49) at various oblique illumination angle as shown in Fig. S1. Fig. 1B (iv-vi) illustrates the fluorescence signal from F-actin of the fixed fibroblast cell at an angle of 22.7°, 60° (HiLo) and 63° (TIRF) respectively. With increased oblique illumination angle, imaging depth of field is also reduced.

##### B. Determination of illumination angle

The scan radius at the back focal plane of the objective is firstly measured by recording the position of the back-reflected beam at the conjugation. Fig. S4A and 4B demonstrated the maximum intensity projection of the reflected beam at various illumination angles at oil-air and oil-water interfaces. The sudden jump in reflected intensity, indicates having reached a critical angle where most of the light was being reflected by the coverslip. The theoretical value of critical angle and maximum angle can be achieved by the objective can be calculated using Eq. 1 and 2, where $\theta_{max}$ =78.98° and $\theta_{critical}$ = 61.4° and 41.2° for oil-air and oil-water interfaces. Those values and the measured radius are then substituted into Eq. 3 for calculating illumination angle θ. The plot of galvo voltage and scan radius is shown in Fig. S4C, while the scan radius corresponding to the illumination angle is shown in Fig. S4D.

$$\theta_{critical} = \arcsin\left(\frac{n_2}{n_1}\right) \quad (1)$$

$$\theta_{maximum} = \arcsin\left(\frac{NA}{n_2}\right) \quad (2)$$

$$r = f_{obj} n_1 sin\theta \quad (3)$$

**C. Quantification of signal to noise ratio (SNR)**

The foundation of interferometric scattering signal ($I_{det}$) lies in three components: namely back-reflection intensity; known as reference field ($I_r = |\overline{E_r}|^2$) pure scattering field ($I_s = |\overline{E_s}|^2$) known as sample field in ROCS [15], and the superposition of reference and scattering ($2E_rE_s\cos\phi$, wherein $\phi = \phi_r - \phi_s$ are the relative phase difference between sample and reference) that refers to interferometric scattering [27, 43].

$$I_{det} \propto |\overline{E_r} + \overline{E_s}|^2 = I_r + I_s + 2E_rE_s cos\phi \quad (4)$$

For nanoscale imaging, scattered intensity ($E_s$) is a few folds (~x) lower than the reference intensity ($E_r$). The superpositioning of the reference and sample amplitude $E_rE_s$ would be imbalanced, resulting in poor interference visibility. To access interferometric scattering, it is crucial to ensure that the superposition intensity $2E_rE_s\cos\phi$ is not being masked by a dominant reference field ($I_r = |\overline{E_r}|^2$).

The gold/polystyrene particles were firstly diluted to desired concentrations using distilled water and dried on a no.1 coverslip. Prior to an imaging session, a drop of distilled water (100 µL) was added to the nanoparticles adhered onto a coverslip via Van Der Waals forces. To calculate the SNR of the nanoparticles at a specific illumination angle, a minimum of 50 regions containing single particles were cropped from the original image, captured by the camera. The peak value and averaged background were used for computing SNR using Eq. 5 and outliers (beyond ± 1.5*std) were removed.

$$SNR = \frac{abs(I_s - I_{bkg})}{I_{bkg}} \quad (5)$$

**D. Fixation and staining of fibroblast cell**

An L929 murine fibroblast cell line was maintained in Dulbecco's modified Eagle's medium containing 10% fetal bovine serum in a humidified 10% CO2 atmosphere at 37 °C. Before each experiment, cells were re-plated onto custom-made polydimethylsiloxane (PDMS) imaging chambers and grown for an additional 1−2 days until a confluency of ~70% coverage was reached. Cells were fixed with 4% paraformaldehyde (Thermo Scientific) in PBS for 15 minutes, followed by three PBS washes and then permeabilized with 0.1% Triton X-100 for 15 minutes. Permeabilized cells were washed with PBS three times and then incubated with Alexa Fluor 488 conjugated phalloidin (Actin Green 488 ReadyProbes, Invitrogen) for 30 minutes to visualize intracellular actin followed by three PBS washes before imaging.

**E. Isolation and preparation of human platelet**

The protocol of isolation human platelets was adapted [4]. Venous whole blood was collected in acid-citrate-dextrose (ACD; 97 mM trisodium citrate, 111 mM glucose, 78 mM citric acid). To obtain washed platelets, whole blood was firstly diluted 1:1 with modified Tyrode's buffer (137 mM NaCl, 2.8 mM KCl, 12 mM $NaHCO_3$, 5.5 mM glucose, 10mM HEPES, pH 6.5) and centrifuged at 70 g for 30 min at room temperature with no brake. The supernatant containing platelet rich plasma (PRP) was diluted 1:3 parts with modified Tyrode's buffer containing 0.1 µg/ml $PGI_2$ (Abcam) and centrifuged at 1250 g for 10 min at room temperature (RT). The pellet was then re-suspended in modified Tyrode's buffer (with adjusted pH 7.4) to achieve a platelet concentration of $5 \times 10^8$ platelets/mL.

**F. Fibrinogen/fibrin coating**

The fibrinogen/fibrin coating protocol is adapted and modified [4]. Coverslips (#1) was firstly immersed in 20% $NHO_3$ for 1 hour and washed with distilled water (ddH2O). The acid-washed coverslips were then immersed in ddH2O for another hour and rinsed again with $ddH_2O$. The cleaned coverslips were then dried on a coverslip rack. Before fibrin/fibrinogen coating, a layer of HMDS was silanised on the coverslip by vapor deposition to create a hydrophobic surface for protein binding. The dried coverslips were plasma cleaned for 4 minutes and then placed in a vacuum desiccator together with a vial containing 1 ml of HMDS solution at room temperature. The desiccator was connected to a vacuum pump in a fume hood to deposit the HMDS vapor for 40 minutes. The effectiveness of HMDS coating is verified by checking the contact angle of the surface.

After the HMDS coating, a customized PDMS chamber (radius 10 mm, depth 6mm) was placed on the coverslip and 200 ul fibrin (300 μg/ml fibrinogen, 12 mg/ml BSA, 2U/ml thrombin, and 1 mM $CaCl_2$ supplemented in modified Tyrode's buffer, pH=7.4) or fibrinogen solution (300 μg/ml fibrinogen and 12 mg/ml BSA supplemented in modified Tyrode's buffer) was loaded into the chamber. These solutions were used immediately after mixing. For control experiments, modified Tyrode's buffer (pH=7.4) is added instead of BSA. The fibrin and fibrinogen coverslips were coated for 30 minutes and 1 hour at room temperature, respectively and then washed with modified Tyrode's buffer (pH=7.4) three times before use.

##### G. Platelet migration

Washed platelets was supplemented in modified Tyrode's buffer (pH=7.4) with 1 mg/ml BSA, 200 μM CaCl2, 4μM U46619 and 10 μM ADP at final concentration of 107 platelets/ml. The imaging chamber was incubated at 37 °C for 40 minutes to allow platelet adhesion and migration before imaging. Note that incubation at 37 °C is recommended but not a determinant factor for migration. Platelet migration is also observed at room temperature but at a much slower speed and shorter trajectory.

##### H. Morphological profiling of single platelet during migration

The processing of quantifying morphological features is illustrated in Fig. S5. The fluorescence and scattering images are firstly overlapped to recognize migrating platelets. The criteria we used to check if a platelet is migrating is to confirm if there is a dark region associated with the platelet. The area of the dark region needs to be larger than half of a platelet size. We then cropped scattering images of the migrating and non-migrating platelets into a small field of views (500×500 pixels) using Fiji (ImageJ). We put them into a different group for shape analysis using a Cell profiler. The total number of platelets being processed is 937, including 427 on fibrin surface (288 platelets with BSA, 139 without BSA) and 510 on fibrinogen surface (318 platelets with BSA, 192 without BSA). To generate binary segmentation of the platelets, the following modules are used in the processing: rescale intensity, identify primary object (platelet, with adaptive threshold, otsu method), identify secondary object (filopodia associated with the identified platelet, adaptive threshold, robust background method). The processed data is reviewed and edited manually if two objects touch each other. Morphological features of the platelet are calculated using "measure object size and shape" module in the Cell profiler. As shown in Fig. S5C, the area is calculated as the product of the number of pixels covered in the object and the actual pixel size of the imaging system (0.025 $\mu m^2$/pixel). The convex area is the area of the convex hull that encloses the object, and the solidity is defined as the object area/convex area. In Fig. S5C, we demonstrated the filopodia rich shape platelet can have a solidity of 0.39-0.7 while the round or half-moon shape platelets is around 0.85-0.94.

#### I. Tracking intensity along lamellipodia during platelet migration

To track the intensity on the lamellipodia of the platelet, a binary mask showing only the edge of the platelet was firstly generated based on the scattering signal using ImageJ (auto local threshold function with Nilblack algorithm and radius = 8). As the membrane ruffles during migration and causes lamellipodia shifts in the lateral position, a stack of binary masks that corresponds to the lateral part of the edge at a different time point is generated to filter the intensity at lamellipodia only. After that, 14 regions on the lamellipodia were manually selected in the binarized image and then converted to ROIs using the BIOP image analysis plugin. The ROIs were then selected on the interference image to quantify mean intensity. Because of varying intensity in scattered signals, the cell outline might appear discontinuous in some frames after binarization and creating multiple small ROIs in one frame, a separate MATLAB script is used to re-calculate mean intensity when multiple ROIs were presented.

#### J. Live endothelial cell culture and imaging

Human microvascular endothelial cell line (HMEC-1, CRL-3243, ATCC) and were maintained in MCDB131 medium (Invitrogen) supplemented 10% heat-inactivated fetal bovine serum (Sigma), 5 U/ml of penicillin, streptomycin and neomycin (Invitrogen), 2 mM of L-glutamine (Invitrogen), 10 ng/mL epidermal growth factor (Invitrogen) and 1 µg/ml of hydrocortisone (Sigma). Primary human microvascular cells derived from the lung (HMVEC, Lonza) were maintained in EGM2-MV2 Bulletkit (Lonza). At 37°C and 5% CO2. For imaging experiments, HMVEC cells were seeded onto glass bottom dishes (ibidi) at a concentration of $2.5 \times 10^4$ cells/mL coated with Poly-L-Lysine (PLL) for 30 minutes and incubated at 37°C for 24 hours before imaging. Cells were stained for 30 minutes with Mitotracker Green FM (100 ng/mL, Invitrogen) prior to imaging and kept on a heat plate (37°C) during imaging.

#### K. Segmentation of mitochondria

Segmentation was performed on fluorescence video using the MATLAB software Mitometer [68] using the 2D analysis function and omitting fission and fusion events. The segmentation masks were exported from Mitometer and thresholded in Fiji to select the mask. The selection was then exported as region of interests (ROI) using the Analyze Particles function. The ROIs were applied to the scattering or interference videos and all signal outside of the ROIs were cleared.

#### Supplementary figures

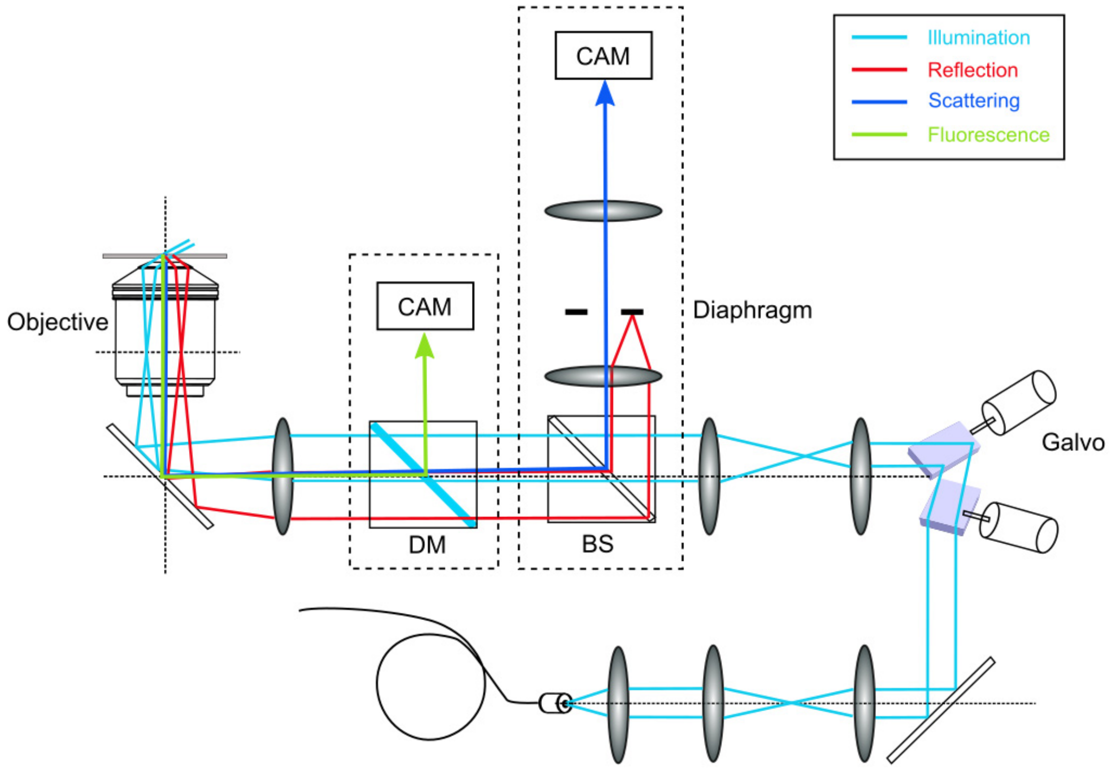


**Fig. S1 Optical setup of combined interferometric, scattering and fluorescence imaging system.** The imaging system is constructed on an inverted microscopy platform (RAMM, ASI) using a single laser source (Stradus 488-150). A two-axis galvo mirror (GVS212/M, Thorlabs) focuses the laser beam onto the back focal plane of the objective lens (Olympus, 60X, NA 1.49, oil immersion), and project radical scanning pattern that allows oblique illumination and azimuthal scanning. The fluorescence and scattering signal are separated using a dichroic mirror (FF505-SDi01-25x36, Semrock) and collected by a CCD (Retiga 4000R, QImaging) and a sCOMS (PCO edge 4.2) respectively.

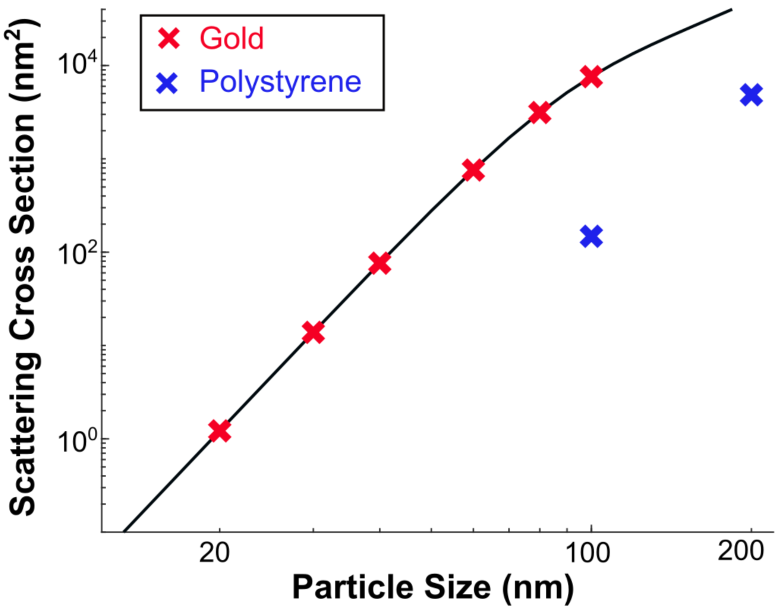


**Fig. S2 Scattering cross section of gold (20 nm, 30 nm, 40nm, 60nm, 80 nm and 100 nm) and polystyrene particles (100 nm and 200 nm), calculating using Mie theory.** The calculation of scattering cross section is based on illumination wavelength of 488 nm, refractive index of the medium and particles to be 1.33, 1.1271+1.8382i (gold) and 1.59 (polystyrene) respectively. The plot is in log-log scale.

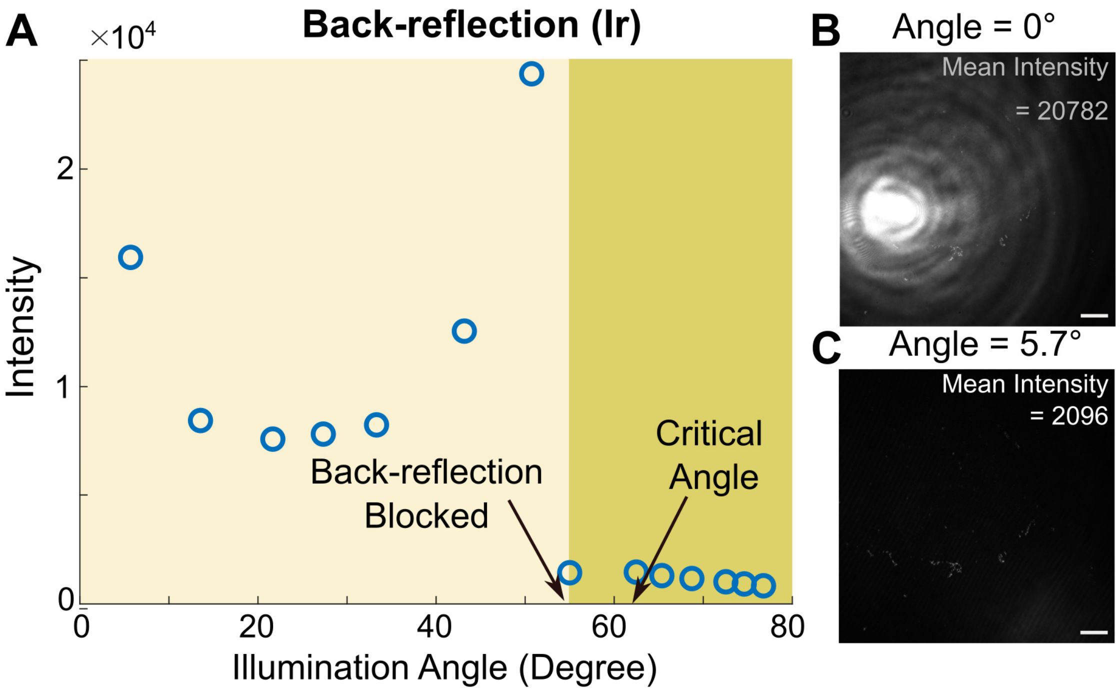


**Fig. S3 Measurement of back-reflection intensity from the coverslip. (A)** Plotting of back-reflection meaning intensity at various illumination angle (5° to 76°). The intensity is measured by capturing reflected light of an empty coverslip covered with water (refractive index = 1.33) at fixed exposure of 30 ms. Example of back-reflection signal and calculated mean intensity with **(B)** on-axis illumination (θ = 0°) and (B) oblique illumination (θ = 5.7°), measured at exposure of 5 ms. Scale bar = 10 μm

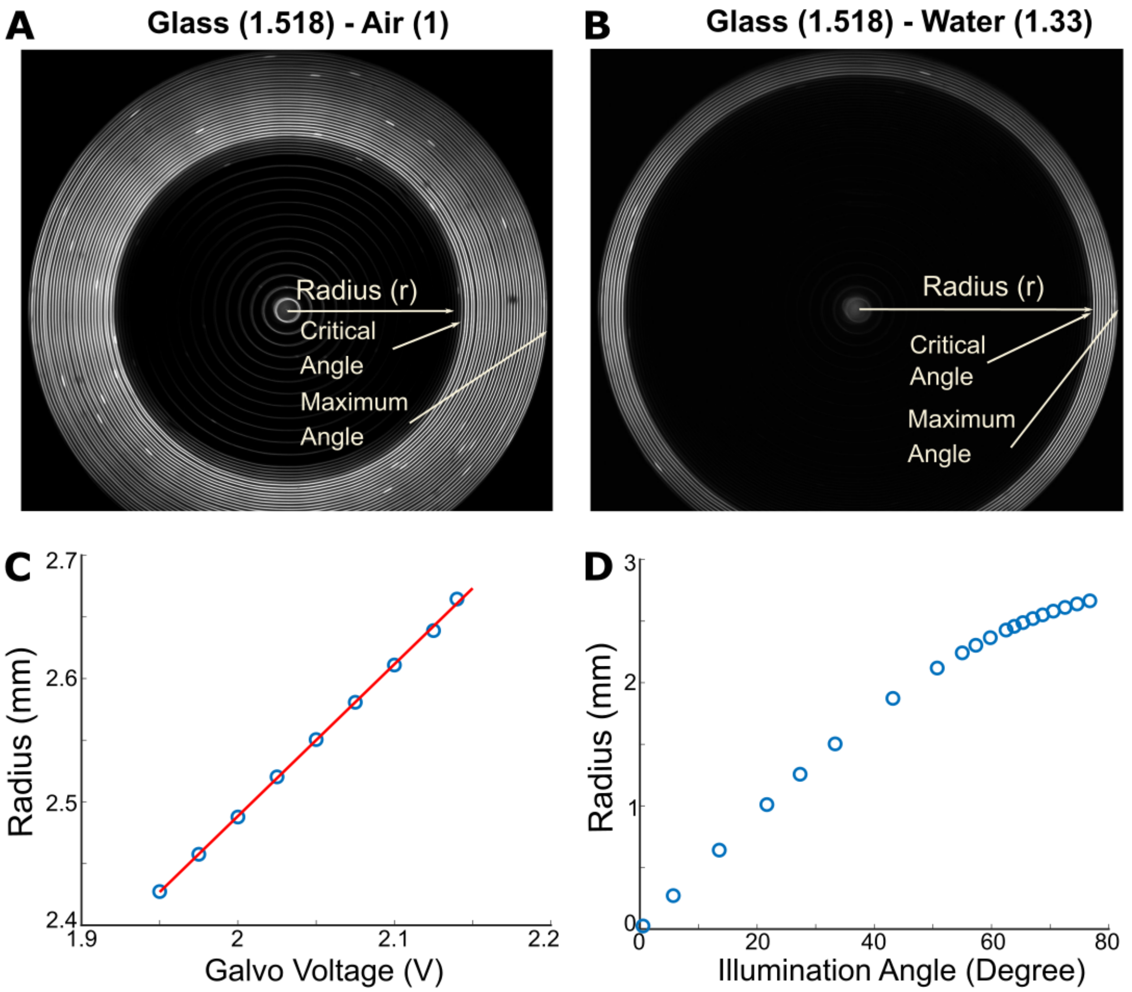


**Fig. S4 Measuring oblique illumination angle.** Recording of the reflected light at back focal plane of two different mediums (A) air (refractive index = 1) and (B) water (refractive index = 1.33) (C) Plotting of measured scanning radius and galvo voltage. Red line shows the fitted linear relation between radius and voltage. (D) Plotting of calculated the illumination angle vs. scanning radius.

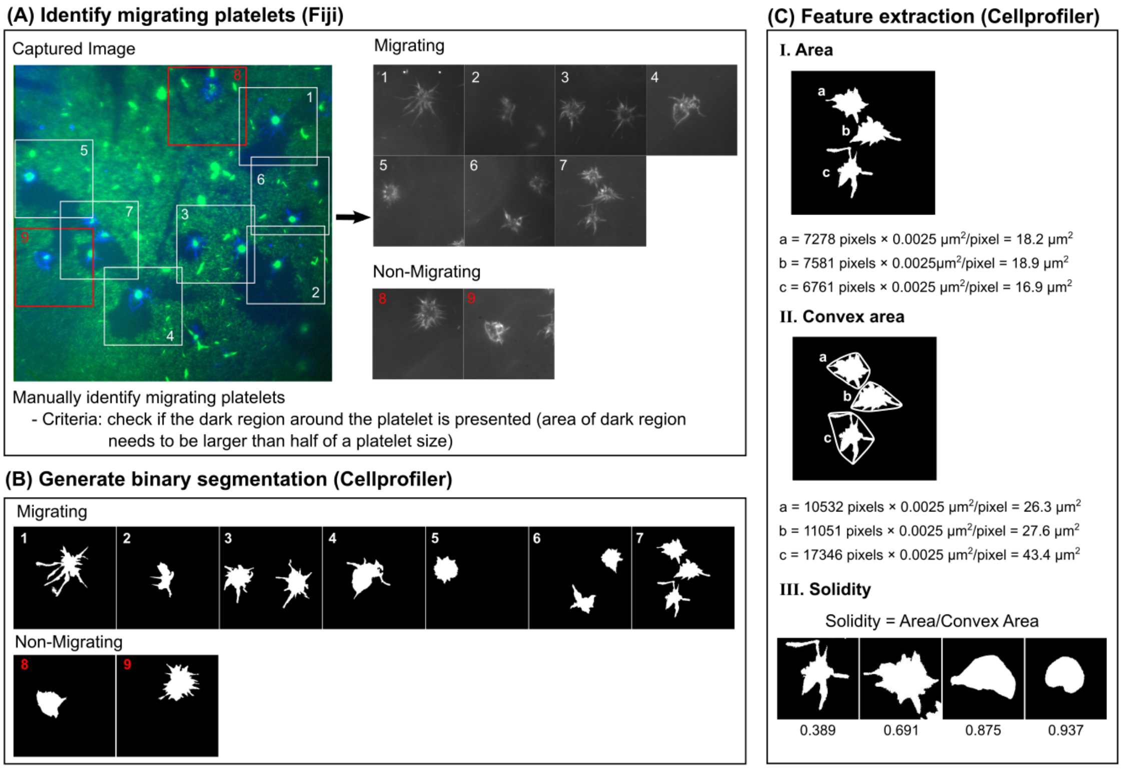


**Fig. S5 Quantifying morphological features of platelets using Fiji (ImageJ) and *Cell profiler*. (A)** The migrating platelets are firstly identified manually using the overlapping image of fluorescence (green) and scattering (blue) signal. The criteria for migration is to check if there is a dark region associated with the platelet and the area of the dark region needs to be larger than half of a platelet size. The migrating and non-migrating platelets are then cropped into small field of views (500×500 pixels) and put into different group for shape analysis. (B) After that, the binary segmentation is generated using cell profiler and the morphological features (area, convex area and solidity) are calculated. (C) These quantifications are illustrated and explained with examples.

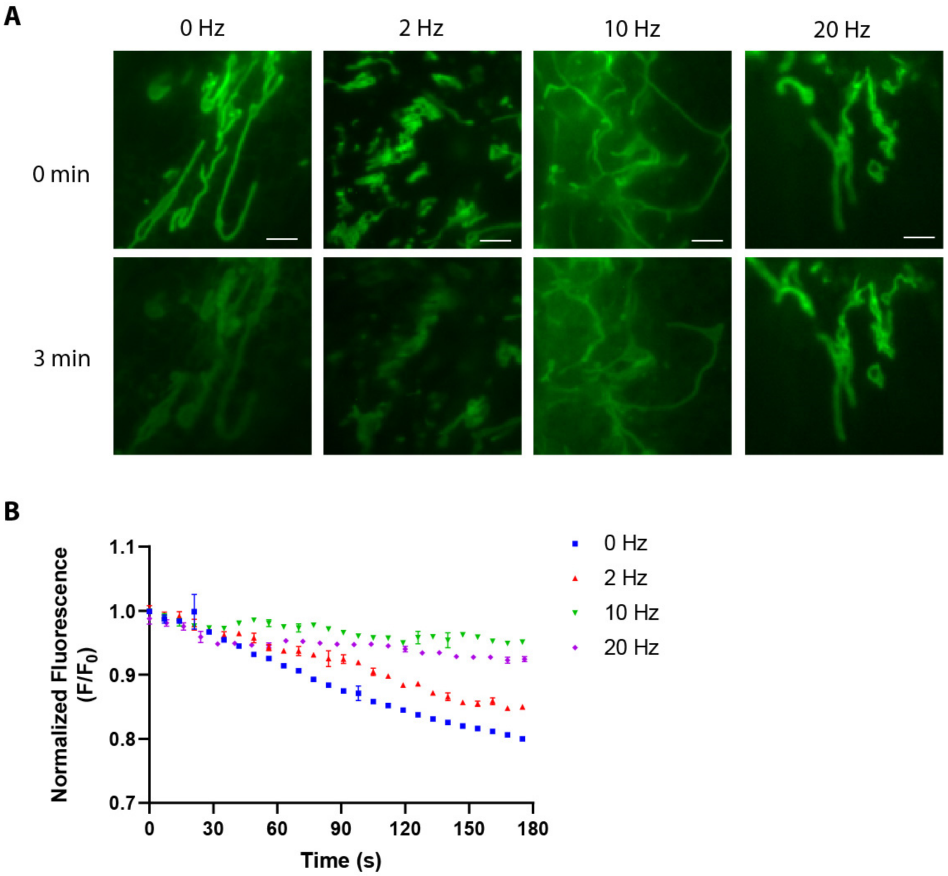


**Fig. S6. Reducing photobleaching with laser pulsing.** (A) Fluorescence imaging of primary human endothelial cells stained with Mitotracker Green before and after 3 min illumination with 0 – 20 Hz laser pulse. (B) Plot of Mitotracker fluorescence in measured in (A), normalized to intensity at start of imaging and plotted over illumination time. Intensity values are the average of 5 frames.

**Movie 1-** Interferometric, DF, BF and TIR ROCS imaging of fixed HMVEC cells and intensity profile plot along a filopodium.

**Movie 2-** Simultaneous recording of both scattering and interference of single platelet spreading on fibrin coated surface

**Movie 3 -** Video recording of leading edge of filopodia of a single platelet being folded into a leading lamellipodium after adhesion on fibrin surface

**Movie 4** –Interference, scattering and fluorescence imaging of a single platelet migrating on fibrin coated surface.

**Movie 5** – Automated tracking of platelet interference images along the boundaries during migrating (highlighted in green) on fibrin surface. The boundaries are segmented using auto local threshold function with Nilblack algorithm in ImageJ.

**Movie 6**– A single sub-region tracking (highlighted in green) on the rear edge of interference images of platelet migrating on fibrin surface using ImageJ.

**Movie 7** – Real time interference, scattering and fluorescence imaging of migrating live HMEC-1 endothelial cells labelled with Mitotracker Green. Mitochondria were segmented using the Mitotracker fluorescence signal. Videos of expanded FOVs show real-time axial and lateral motion of mitochondria and membrane retrograde flow at the cell's leading edge.

**Movie 8** – Interference and fluorescence imaging of a migrating HMEC-1 cell stained with Mitotracker Green. Highlighted sections show interference imaging of membrane retrograde movement at the cell leading edge and fluorescence imaging of mitochondria lateral motion. Expanded video reveals time point when a mitochondrion enters the nucleus.
